# Microbial iCLIP2: Enhanced mapping of RNA-Protein interaction by promoting protein and RNA stability

**DOI:** 10.1101/2024.10.15.618220

**Authors:** Nina Kim Stoffel, Srimeenakshi Sankaranarayanan, Kira Müntjes, Nadine Körtel, Anke Busch, Kathi Zarnack, Julian König, Michael Feldbrügge

**Author notes:** Dr. Michael Feldbrügge, Institute of Microbiology, Cluster of Excellence on Plant Sciences, Collaborative Research Center “Microbial Networking” Heinrich Heine University Düsseldorf, 40204 Düsseldorf, Germany.

## Abstract

The entire RNA lifecycle, spanning from transcription to decay, is intricately regulated by RNA-binding proteins (RBPs). To understand their precise functions, it is crucial to identify direct targets, pinpoint their exact binding sites, and unravel the underlying specificity *in vivo*. Individual-nucleotide resolution UV crosslinking and immunoprecipitation 2 (iCLIP2) is a state-of-the-art technique that enables the identification of RBP binding sites at single-nucleotide resolution. However, in the field of microbiology, optimized iCLIP protocols compared to mammalian systems are lacking. Here, we present the first microbial iCLIP2 approach using the multi-RRM domain protein Rrm4 from the fungus *Ustilago maydis* as an example. Key challenges such as inherently high RNase and protease activity in fungi were addressed by improving mechanical cell disruption and lysis buffer composition. Our modifications increased the yield of crosslink events and improved the identification of Rrm4 binding sites. Thus, we were able to pinpoint that Rrm4 binds the stop codons of nuclear-encoded mRNAs of mitochondrial respiratory complex I, III and V – revealing an intimate link between endosomal mRNA transport and mitochondrial physiology. Thus, our study serves as a paradigm for optimizing iCLIP2 procedures in challenging organisms or tissues under high RNase/ protease conditions.

## Introduction

RNA-binding proteins (RBPs) are ubiquitous and play a pivotal role in determining the physiological state of the cell. Dysfunctional RBPs have been implicated in various human diseases (Gebauer et al., 2021). Hence, it is crucial to investigate RNA-protein interactions *in vivo* with a focus on understanding the interactome, its specificity, and regulation.

Depending on the research question, RNA-protein interactions can be studied from either an RNA-or protein-centric perspective (Hentze et al., 2018, Ramanathan et al., 2019, Hafner et al., 2021). The protein-centric approach facilitates the identification of unknown RNA targets for the RBP of interest and can be further categorized into two major groups. Methods involving chemical modifications of bound target RNAs (e. g. RNA tagging, TRIBE; Qin et al., 2021) and techniques that include protein purification like RIP (RNA immunoprecipitation, Tenenbaum et al., 2000), CLIP (UV crosslinking and immunoprecipitation, Ule et al., 2003), and CRAC (UV crosslinking and analysis of cDNA, Granneman et al., 2009). In particular, iCLIP (individual-nucleotide resolution CLIP, König et al., 2010) stands out due to its high resolution, allowing the identification of precise binding motifs. The advantages of the method inspired the establishment of numerous optimized variants mainly in mammalian cells such as irCLIP (infrared CLIP, Zarnegar et al., 2016), eCLIP (enhanced CLIP, Van Nostrand et al., 2016), and iCLIP2 (Fig. 1A; Buchbender et al., 2020).

**Figure 1.**
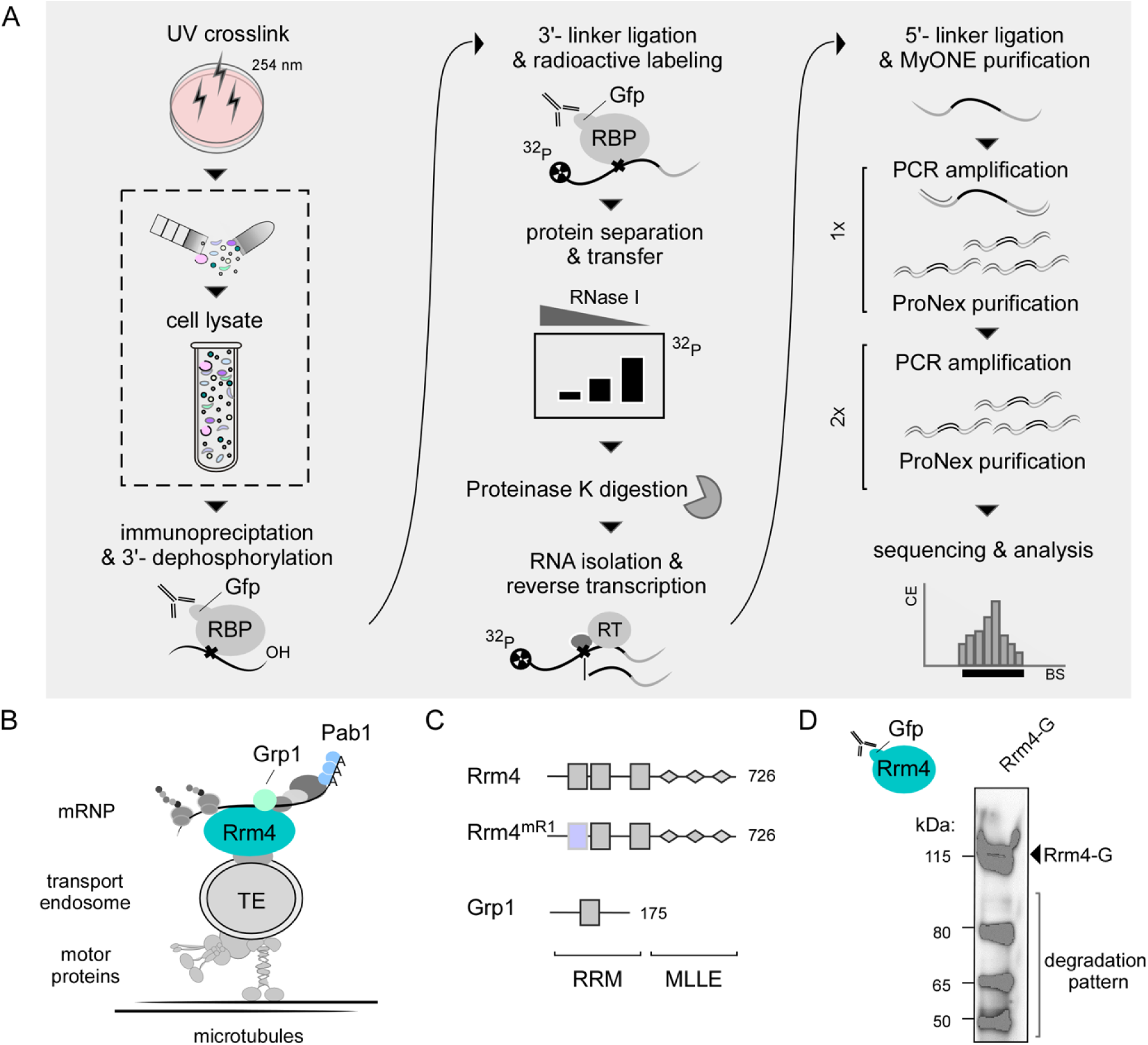
Adaption and improvement of iCLIP2 procedure. **(A)** Diagrammatic overview of iCLIP2 procedure, modified from Buchbender et al., 2020 with a specific emphasis on cell lysis (marked). **(B)** Endosomal mRNA transport model in *U. maydis*. The RBP Rrm4 (RNA Recognition Motif protein 4, petrol) binds directly to cargo mRNAs (black) and co-transports the associated RBPs like Grp1 (Glycine-rich protein 1, mint) or the Poly(A) binding protein (Pab1, blue) in an RNA-dependent manner. **(C)** Schematic overview of the size (in AA) and domain architecture (RRM = RNA recognition motif; MLLE= MademoiseLLE domain) of Rrm4, the amino acid substitutions mutant Rrm4^mR1^ and the RBP Grp1. **(D)** Western blot analysis of immunoprecipitated Rrm4-G (Rrm4-G; 112 kDa).

The iCLIP2 protocol (Fig. 1A) includes the fundamental steps of iCLIP. However, drastic improvements were made to the procedure after cDNA synthesis (Buchbender et al., 2020). This leads to reduced material loss, thereby increasing the library depth. In the past, iCLIP has been adapted for use in plants (Meyer et al., 2017, Lewinski et al., 2024a), archaea (Bathke et al., 2020), as well as in fungi (Olgeiser et al., 2019). Since the protocol involves numerous critical and sensitive steps, establishing improved versions such as iCLIP2 in systems with high protease and RNase activities is nontrivial.

We are investigating endosomal mRNA transport, which is best studied in the model microorganism *Ustilago maydis* and highly polarized cells like neurons (Das et al., 2021, Bourke et al., 2023). In *U. maydis*, RNA recognition motif protein 4 (Rrm4) is the key RBP mediating endosomal RNA transport (Fig. 1B; Müntjes et al., 2021). Components of the transport complex are additional RBPs such as the glycine-rich protein 1 (Grp1; Fig. 1B) and the poly(A)-binding protein 1 (Pab1). Rrm4 has an ELAV-like arrangement of a tandem RNA recognition motif (RRM1,2) and an additionally separated third RRM (RRM3, Fig. 1C). At its C-terminus, it contains a binding platform of three MademoiseLLE domains (MLLE, Fig. 1C) important for endosomal attachment (Devan et al., 2022). To analyze the contribution of the different RRMs of Rrm4, variants with amino acid block mutations in the RNA contact region RNP1 of the respective domains were generated (Becht et al., 2006). Pilot studies revealed that all mutants exhibit strongly reduced RNA binding (Becht et al., 2006). Consequently, a comparative iCLIP2 approach posed several technical difficulties, necessitating further improvements of the method. Firstly, the reduced binding efficiency of the variants diminishes the amount of RNA starting material. Secondly, the high molecular weight of Rrm4 (Fig. 1D) makes immunoprecipitation, gel separation, and blotting challenging. Thirdly, the protein Rrm4 is inherently unstable *in vitro* (Fig. 1D). Fourthly, the method needs to be optimized for high RNase and protease activity common in fungi, leading to protein and RNA degradation upon cell lysis (Cortes-Maldonado et al., 2020). Of note, increased RNase and protease activity is also observed in other systems, such as pancreatic and liver tissue, as well as in plants (Augereau et al., 2016, Hafner et al., 2021). Thus, addressing these challenges with a well-defined lysis protocol that ensures high-quality, non-degraded starting material is essential and constitutes a significant advancement.

## Results

### Advanced lysis techniques for consistent and high-quality iCLIP2 data

A crucial factor for maintaining adequate protein and RNA integrity is the implementation of an efficient lysis protocol. This is particularly important to ensure the quality and depth of an iCLIP2 experiment. As an example, we used Rrm4-G, a functional C-terminal fusion with the green fluorescent protein (Gfp, Becht et al., 2006) and Rrm4^mR1^-G, a mutant with substitution of four amino acids in the RNP1 region of the first RRM domain (Fig. 2A). Aliquots of the same batch (see Materials and methods), i.e., hyphal cells expressing either Rrm4-G or Rrm4^mR1^-G, were used as starting material to analyze four different cell disruption techniques (Fig. 2A, Supplemental Table S1). The approaches differed in terms of equipment, time efficiency, throughput/volume accessibility, temperature conditions, and the separation of mechanical and chemical lysis (dry or wet milling; see Supplemental Table S1 and Materials and methods).

**Figure 2.**
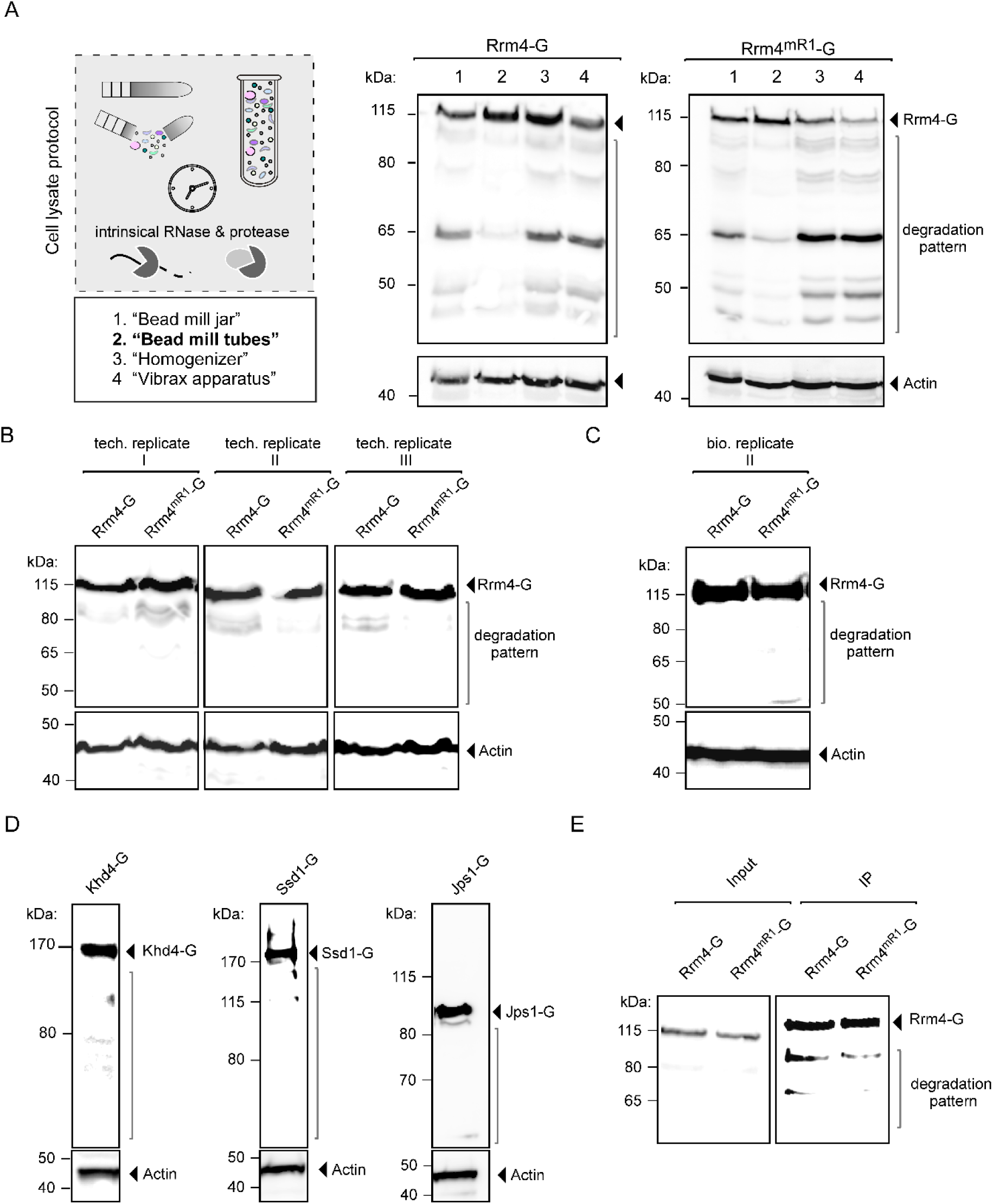
Improved mechanical cell distribution. **(A)** Four techniques to mechanically lyse cells were analyzed (1 = Bead mill jar, 2 = Bead mill tubes, 3 = Homogenizer, 4 = Vibrax apparatus). Start material from the same batches (Rrm4/ Rrm4^mR1^-G) was used. The respective cell lysates were subjected to Western blot analysis (Rrm4/ Rrm4^mR1^-G = 112 kDa, indicated by arrow). The area of protein degradation is marked in dark grey (actin, 42 kDa, served as control). **(B)** Western blot analysis of three technical replicates of technique 2 (Bead mill tubes) are shown. Technical replicates of Rrm4/Rrm4^mR1^-G cell lysates were prepared from the same batches of frozen materials. The full-length protein is indicated by an arrow (112 kDa), and the respective degradation patterns are marked (actin, 42 kDa, served as control). **(C)** Western blot analysis as in B. The respective biological replicates were performed as described previously. **(D)** Western blot analysis of the RBPs Khd4-G (178 kDa), Ssd1-G (180 kDa), and the protein Jps1-G (90 kDa; actin, 42 kDa, served as control). **(E)** Western blot analysis of immunoprecipitation of Rrm4/Rrm4^mR1^-G (IP, 112 kDa, indicated by arrow).

In direct comparison, dry-cryogenic milling („Bead mill tubes“) using a test tube (2 mL) with a steel bead (5 mm diameter) greatly reduced the degradation of Rrm4-G and Rrm4^mR1-^G (Fig. 2A, Supplemental Table S1). In addition, it offers the advantage of processing numerous samples simultaneously in a time-efficient manner (Fig. 2A, Supplemental Table S1). The achieved results are technically (Fig. 2B) and biologically (Fig. 2C; Fig. S1B) highly reproducible, indicating robust and solid cell disruption. Similar results were obtained for additional high-molecular weight RBPs, including Khd4 and Ssd1, as well as the chitinase interactor Jps1 (Fig. 2D; Khd4-G: Sankaranarayanan et al., 2023, Ssd1-G: Bayne et al., 2022, and Jps1-G: Reindl et al., 2020).

Next, we analyzed the compatibility of the improved lysis protocol with the immunoprecipitation approach of iCLIP2 (IP; Fig. 2E) using Rrm4-G and Rrm4^mR1^-G. The input material displayed non-degraded, full-length protein versions (Fig. 2E). As expected, the degradation pattern was slightly increased after immunoprecipitation, attributed to the extended incubation time (see Material and methods). Nevertheless, the experimental setup was proven to be compatible. Thus, the new cell lysis protocol represents a significant improvement in degradation pattern, time efficiency, and simplicity of handling compared to previous protocols (Fig. 1D, 2E).

### Optimizing iCLIP2 buffer conditions for enhanced Protein-RNA complex integrity

Due to the high protease activity in fungal cell extracts, our standard lysis buffer includes 8 M urea to ensure protease denaturation (Devan et al., 2022). Typically, UV crosslinking-based approaches allow stringent highly denaturing conditions like urea and high salt concentrations without affecting the fixed protein-RNA interactions (Challal et al., 2022, Rosenberg et al., 2017). To assess the compatibility of the chaotropic agent urea with our microbial iCLIP2 approach, we verified that the pull-down efficiency remained consistent irrespective of urea concentration for Gfp heterologously expressed in *E. coli* (Supplemental Fig. S1A).

To evaluate the influence of urea, we next analyzed Grp1-G, a small RRM-containing RBP that is highly amenable for iCLIP studies (Meyer et al., 2017, Olgeiser et al., 2019, Lewinski et al., 2024b). Unexpectedly, increasing urea concentrations above 2 M drastically reduced the signal of radiolabeled RNA (Fig. 3A), while the protein abundance remained unchanged (Supplemental Fig. S1C). This could be due to reduced RNA integrity or potentially impacting the radioactive labeling process. Thus, we selected 2 M urea for iCLIP2 experiments with the more challenging protein variants Rrm4-G and Rrm4^mR1^-G. Although the overall radioactive signal decreased when 2 M urea was added to the buffer, the radioactive signal was more concentrated in the region of interest, most likely due to an increase in full-length protein-RNA complexes (above 112 kDa; Fig. 3B-C; Supplemental Fig. S2A-C). Next, we tested the RBP Grp1 as positive control and Gfp as negative control (Supplemental Fig. S2D). As expected, Grp1 showed a strong radioactive signal, whereas Gfp did not (Supplemental Fig. S2D). Thus, by adding urea to our lysis buffer we could reduce unspecific binding to Gfp (Supplemental Fig. S2D-E). In summary, the optimization of the urea concentration led to an increase in full-length protein and thus more starting material was available for the iCLIP2 experiment.

**Figure 3.**
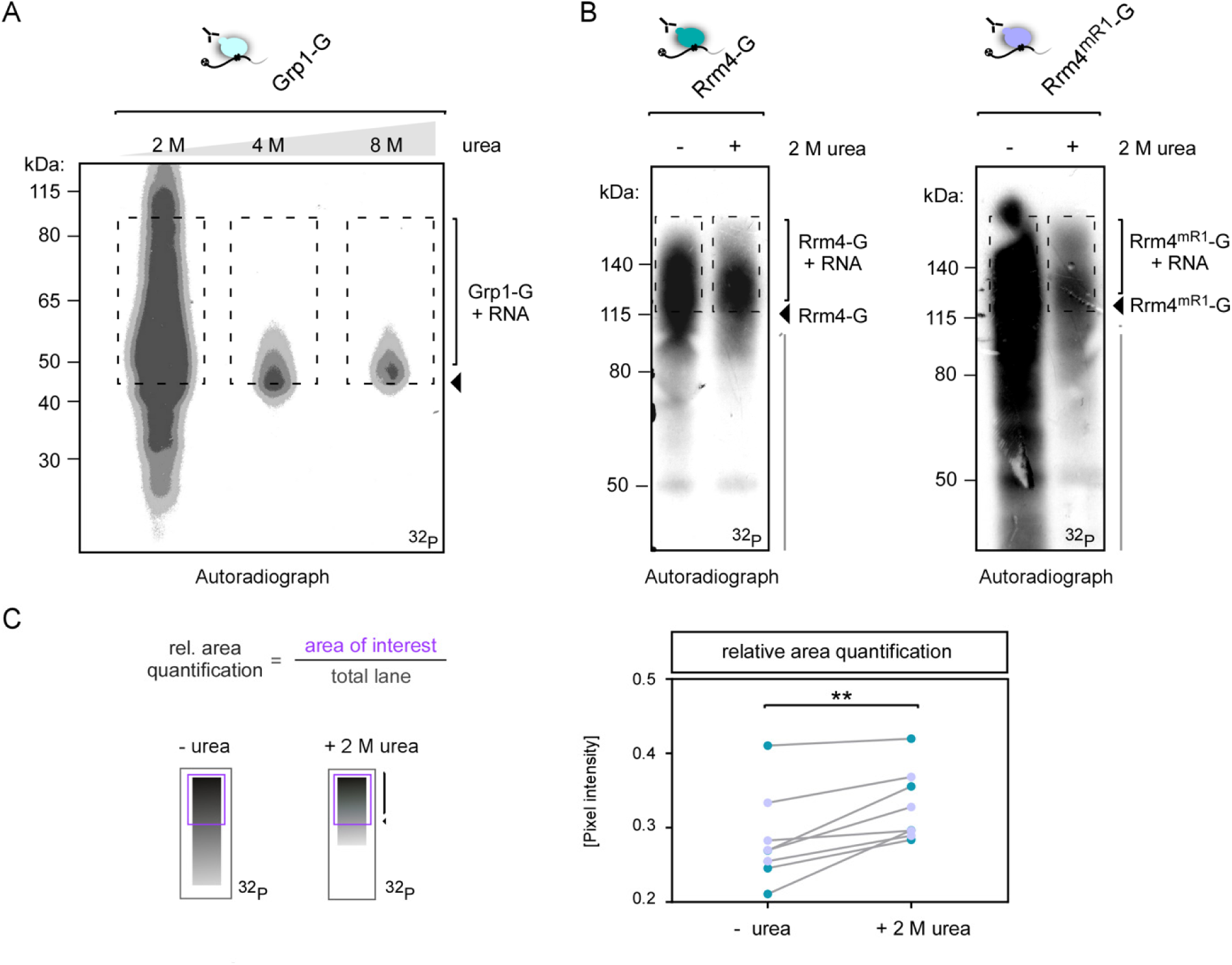
Optimized lysis buffer conditions. **(A)** Autoradiograph of Grp1-G-RNA (area of interest > 45 kDa) complexes, purified with lysis buffer containing 2-8 M urea. **(B)** Autoradiograph of Rrm4/Rrm4^mR1^-G-RNA complexes, immunoprecipitated with or without 2 M urea-containing lysis buffers. The area of interest is indicated (> 112 kDa). The inclusion of 2 M urea in the lysis buffer results in a reduced overall radioactive signal, suggesting higher specificity **(C)** The relative area quantification was determined by the signal intensity of the area of interest (purple) divided by the overall signal of the lane (marked black). Four independent biological replicates of Rrm4 and Rrm4^mR1^-G/RNA-complexes, purified with and without 2 M urea-containing lysis buffer, were quantified (paired T-test; p-value: ** < 0.01).

### Improved cell lysis protocol maximizes RNA integrity

To prepare the sequencing library and to verify the length distribution of PCR products, it is necessary to determine the required minimum number of PCR cycles using a so-called optimization PCR. In contrast to the original protocol that resulted in very short PCR products, we now observed the opposite, i.e. unusually long PCR products (> 250 bp; Fig. 4A) indicative of less RNase activity. We were therefore interested in whether the modified lysis procedure, which reduced protease activity, also contributed to reduced RNase activity. By separating the corresponding area detected within autoradiography (Fig. 4B) into short (S: 115-140 kDa) and long fractions (L: >140 kDa), we confirmed that the cDNA library primarily consisted of longer RNAs (> 250 bp) co-precipitated with Rrm4-G/Rrm4^mR1^-G (Fig. 4B). Thus, our modification of the lysis procedure increased RNA integrity, highlighting the substantial impact of our improvements. Although improved RNA integrity is desirable in principle, too long RNAs can hinder the effective separation of the RNA-protein complexes during gel electrophoresis (Haberman et al., 2017a). Furthermore, libraries prepared from extended RNAs can interfere with standard sequencing read length. Hence, our aim is an optimal range between 150-250 nt.

**Figure 4.**
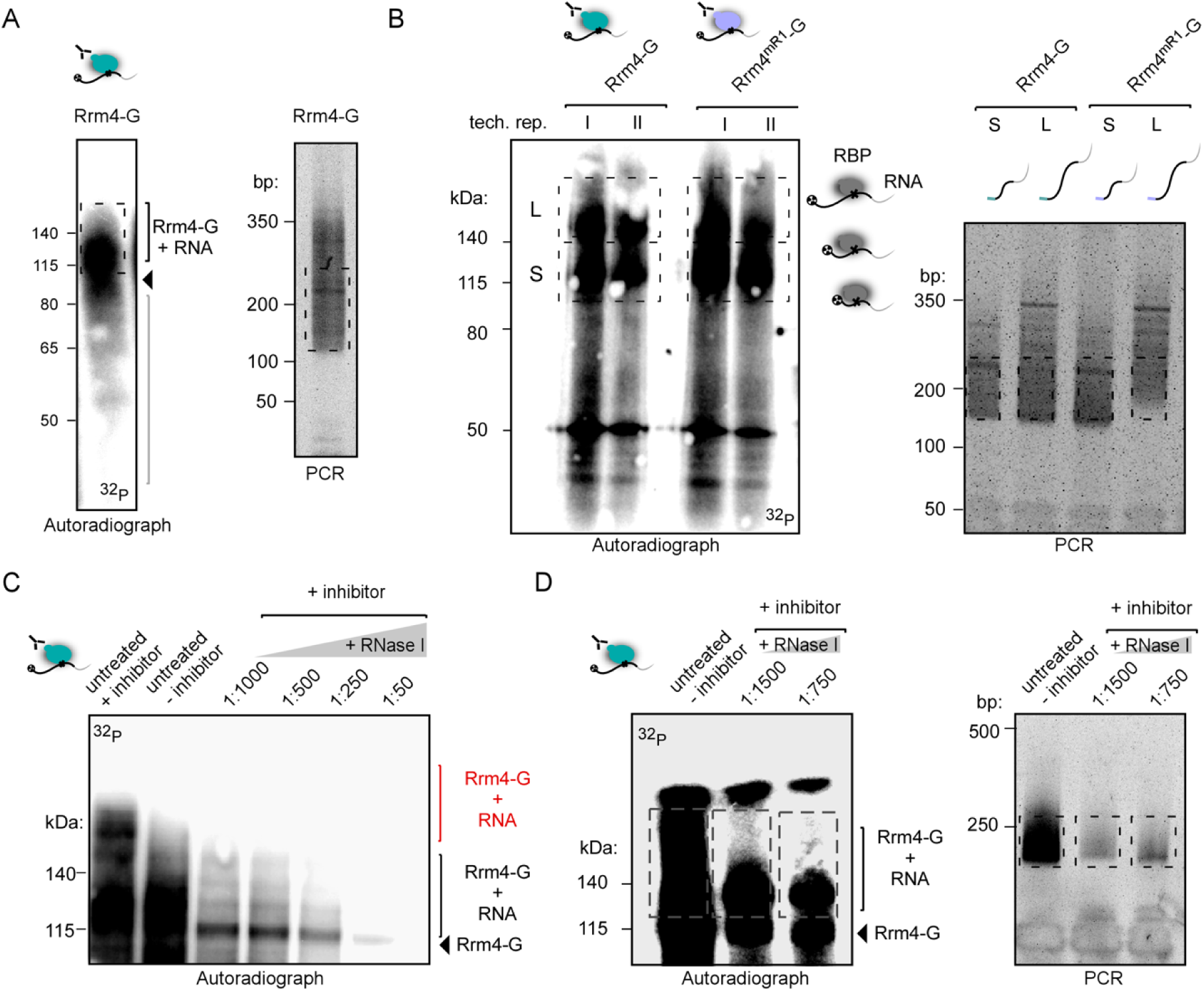
Enhanced RNA integrity. **(A)** Autoradiograph of Rrm4-G-RNA complex purified with 2 M urea. The optimization-PCR showed „unusually long“ > 250 bp cDNA products (optimal range is indicated). **(B)** Autoradiograph of two technical replicates of Rrm4/Rrm4^mR1^-G. The area of interest was segmented into short (marked as S, 112-140 kDa) and long (marked as L, > 140 kDa) RNAs. The respective cDNA libraries (S and L) were used for PCRs. The optimal range of cDNA length is indicated **(C)** Autoradiograph of the RNase I titration series of Rrm4-G-RNA-complexes (> 112 kDa, Rrm4-G full-length band indicated by an arrow, 112 kDa). Indicated in red are Rrm4-G/RNA-complexes exhibiting RNA fragments. The previous protocol procedure (untreated + inhibitor = no RNase I treatment, high amount of RNase inhibitor), internal RNase treatment (untreated, – inhibitor = no RNase inhibitor), and different RNase concentrations (1:1000 – 1:50 RNase I dilution + inhibitor) were analyzed by autoradiography. The inhibitor was added to the cell lysis buffer to reduce internal RNase activity. The respective RNase I (external) treatment was performed after IP within 1x PNK buffer which does not include any RNase inhibitors. **(D)** Autoradiograph and optimization-PCR of Rrm4-G/RNA-complexes under diverse RNase I conditions. The experimental procedure as described in C was repeated with a dilution of 1:750 and 1:1500 RNase I instead, the cDNA library was synthesized and respective optimization-PCRs were performed. PCR products show cDNA sizes in the expected area (marked).

To further improve this step, we titrated the added RNase I within the recommended range of 1:50 to 1:1000 dilutions (Buchbender et al., 2020). Examining the intrinsic RNase activity revealed a strong signal of radioactively labeled RNA even in the absence of RNase inhibitors during lysis and immunoprecipitation steps (untreated, –inhibitor, Fig. 4C). Subsequently, we tested RNase I dilutions of 1:750, 1:1500, and intrinsic RNase activity (untreated, –inhibitor, Fig. 4D). As can be seen in Fig. 4D the intrinsic RNase gave the best results in terms of RNA length. Therefore, we omitted the RNase inhibitor from our lysis buffer for the subsequent steps.

In addition, we also examined the RNase conditions for Grp1-G (Supplemental Fig. S3A-B). Consistent with the previous results for Rrm4-G, Grp1-G co-purified RNAs were rather long in the presence of RNase inhibitors. The smear of the Grp1-G/RNA complex improved under intrinsic RNases (untreated, –inhibitor) and external RNase I titration conditions. Interestingly, for Grp1, a dilution of 1:750 RNase I yielded the best result. This highlights the uniqueness of each RBP (Fig. 4D; Supplemental Fig. S3A-B), underscoring the importance of optimizing the conditions for each RBP of interest individually.

### Reproducibility and enhanced resolution by microbial iCLIP2

To determine the quality and consistency of our microbial iCLIP2 protocol, we further investigated Rrm4-G and Rrm4^mR1^-G (Fig. 5A-C). In both cases, the RBP-RNA complex (above 112 kDa) and cDNA libraries were in the optimal range (about 150-250 bp; Fig. 5B-C). This validates the reproducibility of our procedure, demonstrating a consistent cDNA length without fluctuations attributed to intrinsic RNases (Fig. 5B-C). In addition, we were able to decrease the number of required cycles for the preparative PCR of Rrm4-G compared to our previously achieved Rrm4-G iCLIP experiments (Fig. 5D, our previous iCLIP: 18-22 cycles (Olgeiser et al., 2019); modified iCLIP2: 14-16 cycles). In summary, our protocol is suitable for RBPs with lower binding capacity, as confirmed by Rrm4^mR1^-G.

**Figure 5.**
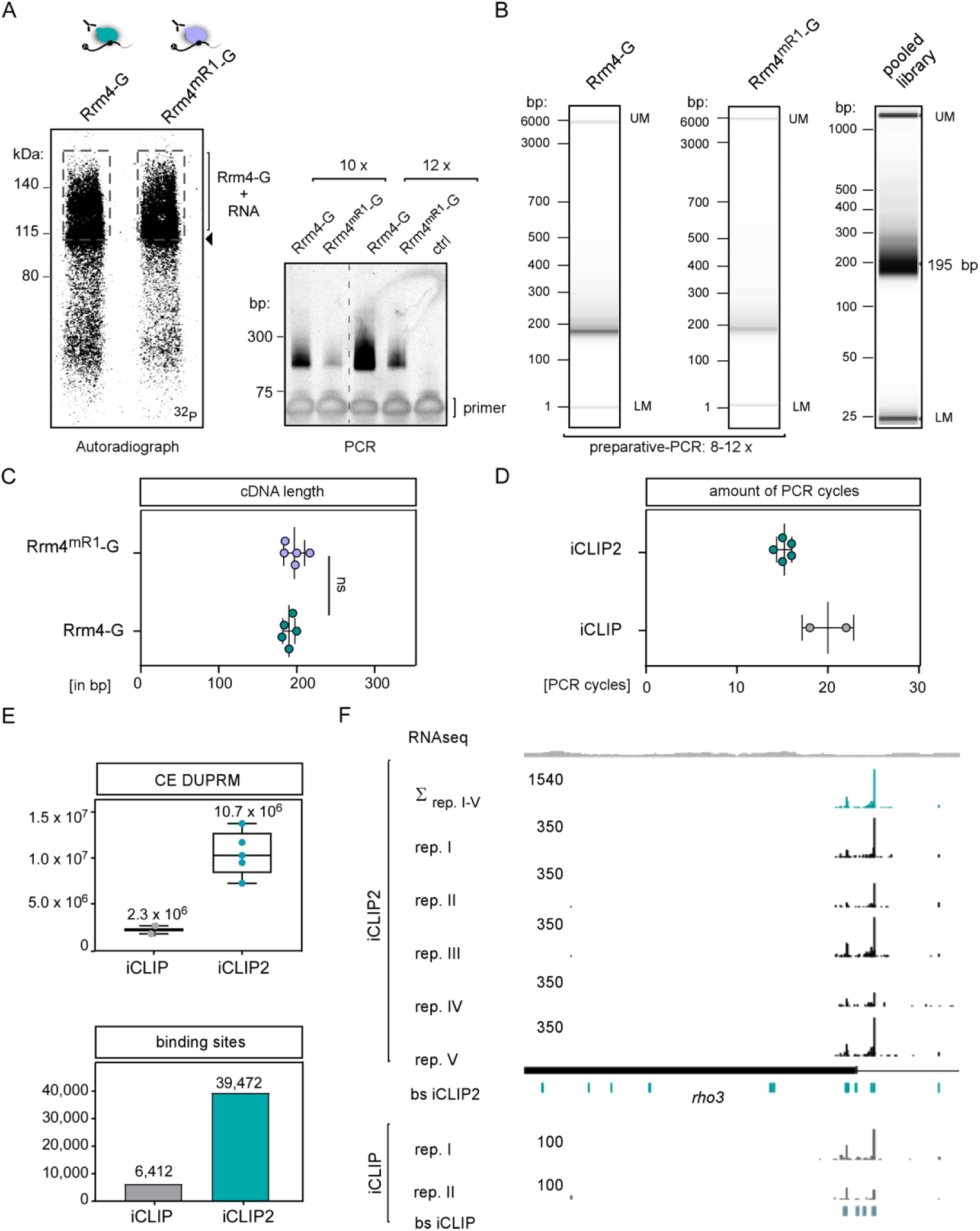
Adaption of improved iCLIP2 protocol. **(A)** Autoradiograph and optimization-PCR of Rrm4/Rrm4^mR1^-G. **(B)** Capillary gel electrophoresis image of the final PCR of Rrm4/Rrm4^mR1^-G libraries before and after pooling. The respective amount of used PCR cycles depends on the RBP (Rrm4 vs. Rrm4^mR1^-G) and the quantity of the respective cDNA library. The pooled library includes additional samples beyond Rrm4-G. **(C)** Reproducible cDNA length based on the mean of n = 5 biological replicates of Rrm4-G/Rrm4^mR1^-G. **(D)** Amount of PCR cycles for Rrm4-G needed for library preparation by iCLIP (18-22 x; n = 2) or iCLIP2 (1. PCR: 6 x; 2. PCR: 8-10 x; in total = 14-16 x; n = 5) procedure. **(E)** Quantified numbers of crosslink events after duplicate removal (CE after DUPRM) of Rrm4-G iCLIP (n = 2) and Rrm4-G iCLIP2 (n = 5). Number of binding sites determined for Rrm4-G in iCLIP (n = 6,412) vs. iCLIP2 (n = 39,472). For both data sets a nucleotide window of 9-nt has been chosen **(F)** Genome browser view of *rho3* mRNA. The numbers represent the upper limit of stack heights for iCLIP(2) crosslink events (CE) and RNAseq reads. The RNAseq coverage of *Wt* filaments is depicted in grey. The five biological Rrm4 replicates of iCLIP2 are depicted in black (mentioned as rep. I-V). The respective merged crosslink events of all five biological replicates and the determined binding sites (bs) are depicted in petrol. The *rho3* ORF is shown as a filled rectangle. The two biological Rrm4 iCLIP replicates (grey, rep. I/II) and the determined bs (light petrol) are depicted underneath (modified from Olgeiser et al., 2019).

Next, we conducted the entire iCLIP2 procedure, analyzing five biological replicates of Rrm4-G, to obtain a more comprehensive understanding of the library quality. Data evaluation and quality control steps indicated that these libraries possessed high quality with the majority of the reads uniquely mapped (Supplemental Fig. S4A). For bioinformatics analysis, we used a slightly modified version of the iCLIP2 data processing workflow described in the original publication (see Material and methods, Busch et al., 2020). We obtained an average of 10.7 million crosslink events after duplicate removal per replicate (n = 5; CE after DUPRM; Fig. 5E, top), a great improvement from the previous data set (n = 2; average of 2.3 million CE after DUPRM, Olgeiser et al., 2019). This suggests a substantial enhancement in the signal-to-noise ratio, thereby improving binding site predictions (Fig. 5E). Also, the majority of binding sites were detected in all five biological replicates (Supplemental Fig. S4B), indicating a high level of reproducibility in the procedure. A comparison of the previous iCLIP (Olgeiser et al., 2019) and the improved iCLIP2 approach revealed that 94% of the previously defined target mRNAs were also detected in iCLIP2, with the identification of an additional 1,757 targets (Supplemental Fig. S4C). As an example of the quality of iCLIP2, the well-characterized Rrm4 binding target mRNA *rho3* is depicted (König et al., 2009; Fig. 5F). All iCLIP2 replicates (rep I-V) show a similar binding (Fig. 5F; Supplemental Fig. S4B). Comparing iCLIP2 and iCLIP reveals a highly reproducible Rrm4 binding pattern, with both datasets showing predominant crosslink events within the 3’ UTR of *rho3* (Fig. 5F). This demonstrates that the redefined protocol and the improved bioinformatic pipeline, including iCLIP data preprocessing (Busch et al., 2020), PureCLIP (Krakau et al., 2017) and BindingSiteFinder (Busch et al., 2020), did not introduce inconsistencies and that, across independent experiments, the RNA interactome remains consistent.

### Rrm4 binds at stop codons of mRNAs encoding mitochondrial proteins of the respiration chain complexes

The previous Rrm4 iCLIP data had exhibited binding at the stop codon of a distinct set of target mRNAs (Fig. 6A; Olgeiser et al., 2019). iCLIP2 showed a similar binding distribution (Fig. 6A). Furthermore, iCLIP2 revealed an expansion in stop codon targets (n = 968; Fig. 6B). The overlap between iCLIP and iCLIP2 (n = 206) demonstrated significant enrichment of the respiratory complex V, as previously reported (Supplemental Fig. S4D; Olgeiser et al., 2019). This new subset (n = 968) significantly enriched for components related to mitochondrial function, particularly within complex I (Fig. 6B). Therefore, we conducted a more in-depth analysis of the *U. maydis* respiratory complex I (Fig. 6C). A comparison of bacterial, fungal, and mammalian complex I components revealed that *U. maydis* comprises 27 nuclear and 7 mitochondrial components, including the evolutionarily conserved core complex N, Q, and P, suggesting a fully functional complex I (Supplemental Table S2). Compared to the human version, *U. maydis* lacks 11 components, which are primarily located in the periphery (Supplemental Table S2), positioning *U. maydis* as an evolutionary intermediate between mammals and *Saccharomyces cerevisi*ae, the latter of which lacks the multimeric complex I.

**Figure 6.**
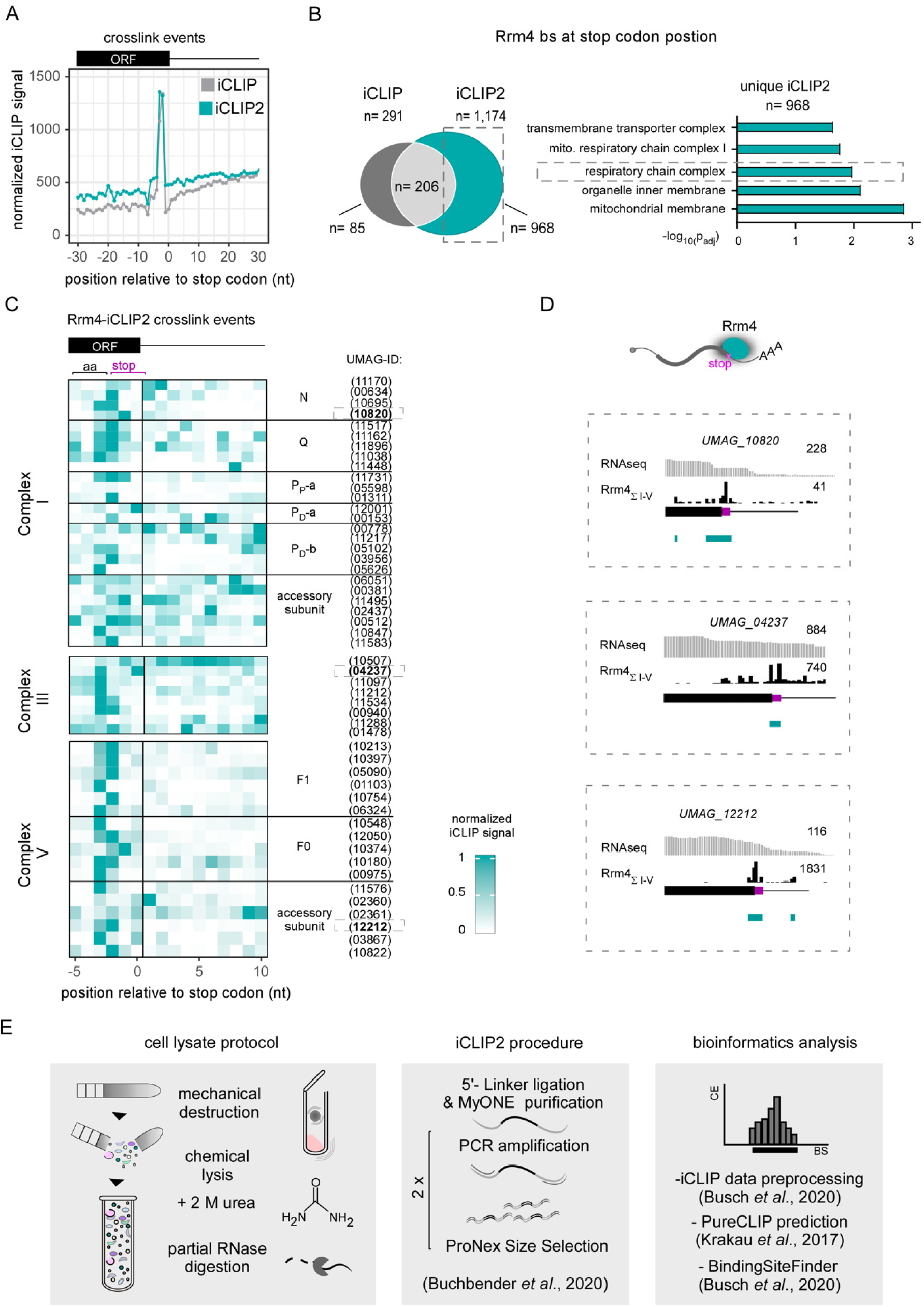
Rrm4 binds at stop codons of nuclear-encoded mRNAs of the respiratory chain complex I, III and V. **(A)** Metaprofile of Rrm4 iCLIP (grey) and iCLIP2 (petrol) normalized signals relative to stop codon **(B)** Venn diagram showing the number of Rrm4 binding sites at the stop codon position as detected by iCLIP and iCLIP2 (dark grey: uniquely found in iCLIP n = 85; light grey: shared targets n = 206; petrol: uniquely found in iCLIP2 n = 968). Selected bar chart from GO-term analysis of stop codon binding targets uniquely determined within the iCLIP2 dataset. **(C)** Heatmap of Rrm4 iCLIP2 crosslink events in a window (–5-nt to 10-nt) around the stop codon position (0 is defined as the last nucleotide of the stop codon). The amino acid codon preceding the stop codon is indicated (aa). The core complex subunits N, Q, and P, as well as the accessory subunits of respiratory complex I, are indicated. Complex III and the subunits of complex V (F_1_ and F_0_) and their accessory subunits are also indicated. The respective identifiers (UMAG_ID) of the individual proteins are listed alongside. **(D)** Genome browser views of *nuo2* (UMAG_10820, Comp I), *qcr7* (UMAG_04237, Comp III), and *atp10* (UMAG_12212, Comp V) mRNAs. The predicted binding sites are depicted in petrol and the stop codon is highlighted in magenta **(E)** Schematic overview of the optimized microbial iCLIP2 procedure. Lysis protocol was improved by the adaption of dry-cryogenic milling, buffer optimization by adding the chaotropic detergents urea (2 M), and internal RNase digestion. The adaptation of the advanced iCLIP2 procedure and defined bioinformatics analysis, including data preprocessing, PureCLIP and BindingSiteFinder (described in Material and methods).

Analyzing transcriptome-wide Rrm4 binding revealed an interesting pattern for mRNAs encoding mitochondrial respiratory complexes. (Fig. 6C). Binding sites at the stop codon position were evident for complexes I, III, and V of the mitochondrial respiratory chain complex within our iCLIP2 dataset (Fig. 6C). Interestingly, for complex I, we found a prominent binding pattern within core subunits N and Q (Fig. 6C). As representative examples of the mitochondrial respiratory complexes, *nuo2* (UMAG_10820, Comp I; Fig. 6D), *qcr7* (UMAG_04237, Comp III; Fig. 6D), and *atp10* (UMAG_12212, Comp V; Fig. 6D) are depicted.

In summary, the enhanced resolution achieved with the improved iCLIP2 protocol has revealed a stronger connection between Rrm4 and nuclear-encoded mRNAs, specifically those encoding components of the respiratory complexes, with an emphasis on stop codon binding (Fig. 6A-D). Rrm4 potentially regulates the transport and translation of these mRNAs, thereby influencing mitochondrial energy production. However, it is important to note that our understanding of this biological mechanism is currently incomplete, and further experiments will be needed.

## Discussion

### Establishing a tailor-made protocol for the study of microbial RBPs

To gain a deeper understanding of RNA biology in microorganisms, we aimed to adopt iCLIP2 (Fig. 6E; Buchbender et al., 2020). To overcome the challenge of high intrinsic RNase and protease activities in our system, a crucial step was the implementation of a new cell disruption protocol involving dry-cryogenic bead-milling for cell lysis. Besides microbes, the method has proven applicability in plants (Box et al., 2011) and mammalian tissues (LaCava et al., 2016, Dou et al., 2020). Its efficiency in terms of time, suitability for high throughput, and ability to enhance RNA integrity, particularly under conditions of high RNase activity such as in pancreatic tissues (Li et al., 2009, Chougoni and Grossman, 2020, Munro et al., 2023) make it a versatile and valuable method.

Our refined lysis buffer, including 2 M urea, resulted in more precisely detected radiolabeled RBP-RNA complexes by improving protein stability. In addition, urea can reduce unspecific binding (Huppertz et al., 2014, Rosenberg et al., 2017, Lewinski et al., 2024a) and denature RNases (Almarza et al., 2009). By the reduction of RNase activity, the RNA integrity increased. Consequently, RNase titration became crucial, and within our setup, internal RNases have provided the best results for Rrm4. Contrastingly, for the RBP Grp1, an external RNase I treatment has proven to be the most effective. This highlights the significance of optimizing the protocol based on the RBP used. It is important to note that relying on intrinsic RNase activity is generally not recommended due to the potential bias it can introduce through cleavage (Haberman et al., 2017b). However, we have not identified such a bias so far.

### Applying an optimized protocol to study microbial RBPs

Due to the improved protocol (Fig. 6E), we achieved a deeper resolution, resulting in the identification of additional binding sites and targets of Rrm4. Hence, we identified Rrm4 binding sites at the stop codons of nuclear-encoded mRNAs for the respiratory chain complexes I, III, and V. Interestingly, our findings further strengthen the connection between vesicle-mediated mRNA transport and mitochondria (Müntjes et al., 2021), a relationship that has also been observed in neurons of *Xenopus laevis* and human (Cioni et al., 2019; De Pace et al., 2024) as well as HEK293 cells (Schuhmacher et al., 2023). However, whether Rrm4 plays a role in the translational regulation of these targets is not known so far. Overall, we gained a more extensive RNA interactome of Rrm4 by iCLIP2. This observation supports the idea that endosomes act as a universal mRNA transporter, contributing to the overall distribution of mRNAs (Olgeiser et al., 2019).

iCLIP studies are rare in organisms like plants and fungi or tissues like pancreatic, which exhibit high RNase/protease activity (Alvelos et al., 2021, Arribas-Hernandez et al., 2021). Generally, the realm of RNA-protein interactions in the microbial kingdom, particularly *in vivo* with high resolution, remains relatively underexplored. In *S. cerevisiae*, one study has utilized the iCLIP/miCLIP protocol (Varier et al., 2022). Most other studies have employed the CRAC methodology (Granneman et al., 2009, Challal et al., 2022, Esteban-Serna et al., 2023), which includes a (tandem) affinity-purification step under denatured conditions and differs in cDNA library preparation. We recommend the optimized iCLIP2 procedure, as it yields highly reproducible cross-link events with excellent library depth.

## Conclusions

In essence, the refined protocol has established a solid framework for successfully implementing a comparative microbial iCLIP2 approach of Rrm4 and respective RRM variants. We are optimistic that our enhanced procedure, specifically designed for biological samples with elevated RNase/protease activity within and beyond microbiology, can yield substantial advantages. Additionally, we envision that our improved lysis procedure is easily transferable to other RNA biochemistry methods, including CLIP, CRAC, RIP, and more. Thus, our study might serve as a paradigm for others.

## Material and methods

### Cultivation of *Ustilago maydis*

The laboratory strain AB33 from *Ustilago maydis* was used to synchronize filamentous growth by changing the nitrogen source (Brachmann et al., 2001). The cultures (OD_600_ = 0.5) were grown in complete medium (CM) supplemented with 1% glucose at 28°C with agitation at 200 rpm (described in detail in Devan et al., 2022). Filamentous growth was induced by shifting the cells to nitrate minimal (NM) medium supplemented with 1% glucose at 28°C, 200 rpm for 6 hours post-induction (h.p.i; Brachmann et al., 2001). The filaments were harvested by centrifugation (7,150 *g*, 10 min, 4°C) and washed with 5 mL PBS (Phosphate-Buffered saline pH 7.4; 137 mM NaCl, 2.7 mM KCl, 10 mM Na_2_HPO_4_, and 1.8 mM KH_2_PO_4_).

### Cell harvesting and lysis

Four different techniques were compared (Fig. 2A).

#### Technique 1: „Bead mill jar“

The cell pellet (50 mL cell culture) was resuspended in 2 mL lysis buffer, transferred into “steel jar” flash-frozen using liquid nitrogen (Supplemental Table S1), and stored at – 80°C (Olgeiser et al., 2019). Two steel beads (12 mm diameter) were added to each jar. Subsequently, the filled steel jars were subjected to three rounds of shaking at 30 Hz for 10 minutes (min) each (RetschMill M400), with intermittent refreezing steps in liquid nitrogen. The frozen lysate was thawed for approximately one hour at 4°C and transferred to a 2 mL reaction tube and centrifuged at 16,200 *g*, 4°C, 10 min. The resulting supernatant was utilized for subsequent steps or flash-frozen as needed. Overall, the experiment is very time-consuming and not well-suited for processing multiple samples simultaneously or handling a large sample volume, primarily because only two steel cups can be run at the same time. To conduct a complete iCLIP(2) approach, each sample typically requires a minimum of 150 mL of culture. For two different samples, this means a total of six jars, 100 min of shaking time, an additional hour for thawing, and the centrifugation step. The entire procedure typically takes approximately 3.5 hours.

#### Technique 2: „Bead mill tubes“

The cell pellet (50 mL culture) was resuspended in 2 mL PBS, split, and transferred into 2 mL reaction tubes (each 1 mL). The tubes were subsequently centrifuged at 16,200 *g*, 4°C, for 5 min, and the supernatant was completely removed. The tubes were placed in a floating holder and flash-frozen in liquid nitrogen (Supplemental Table S1). The tubes can be stored at – 80 °C until use. To each tube, one steel bead (5 mm diameter) was added and placed into a pre-cooled TissueLyser Adapter (Qiagen). The samples underwent two rounds of cell lysis, each lasting for 30 seconds at 30 Hz, with a liquid nitrogen cooling step in between., The resulting powder was reconstituted using 1 mL of lysis buffer. Cellular debris was separated by centrifugation at 16,200 *g* for 10 min at 4°C. The supernatant was carefully transferred, and the samples were pooled again into a fresh pre-cooled tube. An additional centrifugation step was performed for 5 min. The resulting cell lysate can be stored at – 80°C. Utilizing the TissueLyser Adapter (Qiagen) in the RetschMill (M400), up to 48 of the 2 mL reaction tubes (24 tubes per adapter) can be accommodated simultaneously. For optimal thawing, tubes were placed in a Vibrax (Ika-Vibrax-Vxr, Electronic), and resuspension was achieved through gentle mixing for 5 min, < 500 rpm at 4°C. In contrast to strategy 1, this method offers higher throughput and is more time-efficient (1 – 1.5 h).

#### Technique 3: „Homogenizer“

An alternative common method for disrupting difficult cell tissues involves the use of a homogenizer (Precellys 24, Peqlab, Berlin). In this approach, the cell pellet from a 50 mL culture was resuspended in 2 mL of lysis buffer and transferred into 2 mL reaction tubes. A spoon of 0.5 mm glass beads was added, and the homogenization was carried out using a Precellys 24 homogenizer for 1 min at full speed under liquid nitrogen conditions (Supplemental Table S1). The suspension was thawed on ice and centrifuged at 16,200 *g* for 10 min at 4°C to separate the cell debris. The resulting supernatant was carefully transferred into a new tube, and the centrifugation step was repeated for an additional 5 min.

#### Technique 4: „Vibrax technique“

As an additional option, we tested the „Vibrax technique“ (Schmitz et al., 2019). In this method, a 50 mL culture pellet was resuspended in 650 µl of buffer and transferred to a new 2 mL reaction tube containing 300 mg of 0.5 mm glass beads (Supplemental Table S1). The suspension was flash-frozen and subsequently thawed by shaking on a Vibrax apparatus (Ika-Vibrax-Vxr, Electronic) at 2,000 rpm for 30 min at 4°C. The samples were spun at 16,200 *g*, 10 min, and 4°C. The complete supernatant was transferred into a fresh reaction tube and subjected to an additional spin (16,200 *g*, 5 min, at 4°C). The resulting cell lysate can be stored (– 80°C) or directly used for further experiments.

### Western blot analysis

The protein concentrations of cell lysates were determined by Pierce^TM^ BCA assay (Thermo Fisher; 23225) or standard Bradford assay (BioRad, 1976). Samples were adjusted to similar concentrations (1 mg/mL) in NuPAGE LDS sample buffer containing DTT (Thermo Fisher, NP0007; DTT-Dithiothreitol) was added, according to the manufacturer’s instructions. The samples were loaded onto a 4-12% NuPAGE™ Bis-Tris gel (Thermo Fisher, NP0321) and run for 1.5 hours, 180 V. The gel was wet-blotted onto a nitrocellulose membrane (Amersham Protan) using 1x NuPAGE™ Transfer buffer + containing 15% Methanol (Thermo Fisher; NP00061) for 1.5 hours at 30 V. For Western blot detection, α-Gfp (1:2000 dilution; Roche, 11814460001) and α-Actin (1:2000, MP Biomedicals 0869100) from mouse were used as primary antibodies, while α-mouse HRP conjugate (1: 5000 dilution; Promega; W4021) was used as the secondary antibody. All antibodies were dissolved in 1x TBS-T including 5% milk powder. Detailed descriptions of general western blot performance are described in Devan et al., 2022. For detection, the Image Analyzer LAS400 (GE Healthcare) and ECL^TM^ Prime (Amersham, RPN2232) were used, according to the manufacturer’s instructions.

### Microbial iCLIP2 procedure

For the iCLIP2 experiment, 150 mL of filamentous cell culture (OD_600_ = 0.5; 6 h.p.i) was used. The experiment was carried out in a 4°C cold room using material and buffers cooled with ice. The culture was divided into three 50 mL tubes and harvested by centrifugation (7,150 *g*, 10 min, 4°C). The pellets were washed with 5 mL of PBS (Phosphate-Buffered saline pH 7.4; 137 mM NaCl, 2.7 mM KCl, 10 mM Na_2_HPO_4_, and 1.8 mM KH_2_PO_4_) and centrifuged again (7,150 *g*, 10 min, 4°C). The final cell pellets were resuspended in 5 mL of PBS, transferred into a Petri dish (100 x 100 x 20 mm, Sarstedt: 82.9923.422), and kept on ice. The resuspended cells were UV irradiated (254 nm, 1x 200 mJ/cm^2^, Biolink UV-Crosslinker, Vilber-Lourmat) and pooled in a 50 mL tube. Cells were harvested by centrifugation (7,150 *g*, 10 min, 4°C) and resuspended with 13 mL PBS, evenly divided into six 2 mL reaction tubes (≈ 25 mL culture per tube), and disrupted as described before (see „*Bead mill tube*“). The cryogenic cell powder was resuspended in 1 mL lysis buffer (50 mM Tris–HCl, pH 7.4, 100 mM NaCl, 1% IGEPAL CA-630, 0.1% SDS, 0.5% sodium deoxycholate, 2 M urea, 2× Complete protease inhibitor EDTA-free [Merck, 11873580001], 1 mM DTT, 1 mM PMSF [+ 5 µl SUPERase-In, Thermo Fisher AM2696]) per 2 mL reaction tube. The homogenate was centrifuged at 16,200 *g* for 10 min at 4°C and the supernatant was transferred to a new tube and centrifuged (16,200 *g*, 5 min, 4°C). The supernatants were pooled into 50 mL tubes. Protein concentrations were determined using Pierce™ BCA Protein Assay Kits (Thermo Fisher, 23225) and adjusted to match the sample with the lowest concentration (aiming for ≈ 2 mg/mL). Cell debris-free supernatants were distributed into three 2 mL tubes, each containing 20 µL pre-washed GFP-Trap beads (Chromotek; gtma-20). Beads were pre-washed twice with lysis buffer without inhibitors using a magnetic separation rack. Immunoprecipitation was conducted for 1 hour at 4°C with continuous rotation. The beads were pooled in a 2 mL reaction tube and washed four times with 1 mL high-salt buffer (50 mM Tris–HCl, pH 7.4, 1 M NaCl, 1 mM EDTA, 1% IGEPAL CA-630, 0.1% SDS, 0.5% sodium deoxycholate). This was followed by two washes with PNK buffer (20 mM Tris–HCl, pH 7.4, 10 mM Mg_2_Cl, 0.2% Tween-20). Finally, the beads were transferred into a fresh 1.5 mL reaction tube and washed three more times with PNK buffer. The rest of the iCLIP2 procedure was adopted from the original iCLIP2 protocol including information on oligonucleotide design (Buchbender et al., 2020). Western blot analysis was performed as described before (see above). The membrane was washed with PBS, covered with a transparent plastic bag and exposed to a BAS Storage Phosphor Screen (GE Healthcare) for approximately 1 hour or overnight at – 20°C depending on the signal strength. The screen was visualized by a Typhoon^TM^ laser scanner (Cytiva), according to the manufacturer’s instructions. The iCLIP2 libraries were sequenced on an Illumina NextSeq 500 sequencing machine to generate 92-nt single-end reads, which included a 6-nt sample barcode as well as 5+4-nt unique molecular identifiers (UMIs). A more detailed protocol and additional advice are available upon request.

## RNase I treatment

An optional RNase I (Life Technologies, AM2295) treatment was performed after the immunoprecipitation. Various dilutions (ranging from 1:50 to 1:1500) were prepared in 1x PNK buffer (20 mM Tris–HCl, pH 7.4, 10 mM Mg2Cl, 0.2% Tween-20) and added to the beads. The beads were incubated for 3 min, 37°C at 1000 rpm, and subsequently, washed with cold High-salt buffer and handled on ice.

## Processing of iCLIP2 reads and binding site definition

The processing of iCLIP2 reads and the definition of binding sites was conducted mostly as previously described (Busch et al., 2020). In brief, initial quality controls were performed using FastQC (http://www.bioinformatics.babraham.ac.uk/projects/fastqc/). Reads with a Phred quality score below 10 in the barcode or UMI regions were discarded. Remaining reads were de-multiplexed based on the sample barcode, which is found on positions 6 to 11, using Flexbar ((Dodt et al., 2012), version 3.4.0). Barcode and UMI regions as well as adapter sequences were trimmed from read ends using Flexbar with a minimal overlap of 1-nt between the read and the adapter, and UMIs were added to the read names. Only reads with a minimum length of 15-nt were retained for subsequent analysis. Trimmed reads were mapped to the *Ustilago maydis* genome (assembly version Umaydis521_2.0) and its annotation based on Ensembl release 51 (Yates et al., 2022) using STAR ((Dobin et al., 2013), version 2.7.3a). During mapping, up to 4% mismatched bases were allowed without soft-clipping on the 5’end. Uniquely mapped reads were processed to remove PCR duplicates using UMI-tools ((Smith et al., 2017), version 1.0.0). Crosslinked nucleotides were extracted using the BEDTools suite ((Quinlan and Hall, 2010), version 2.27.1) as described in Chapter 4.2 of Busch et al., 2020.

Processed reads from all five replicates were merged into a single dataset, and subjected to peak calling using PureCLIP ((Krakau et al., 2017), version 1.3.1) with default parameters. The resulting crosslink sites were used to define binding sites with the R/Bioconductor package BindingSiteFinder (10.18129/B9.bioc.BindingSiteFinder, version 1.7.9). Pre-filtering steps were implemented to retain only the most significant sites. First, crosslink sites ranked in the lowest 5% by PureCLIP score—a measure of binding affinity strength provided by PureCLIP— were excluded. Second, to account for gene expression levels and to improve the binding site quality, crosslink sites with the lowest 10% PureCLIP score in each gene were filtered out. For defining the binding sites, processed crosslink sites within a distance closer than 8-nt were merged, requiring at least 2 out of 9 sites to be covered by crosslink events. The centers of the binding sites were iteratively positioned at crosslink events with the highest PureCLIP score and extended by 4-nt on either side, resulting in binding sites with 9-nt width. Furthermore, binding sites were deemed reproducible if they harbored more crosslink events than the 5th percentile of the crosslink distribution across all binding sites, with at least 2 crosslink events per binding site, observed in four out of five biological replicates.

To assign gene and transcript regions to the binding sites, we utilized the gene annotation file version *U. maydis* 521_2.0.41. Genes in the *U. maydis* genome were manually extended by 300-nt on each side to encompass potential 5’ UTR and 3’ UTR regions (Sankaranarayanan et al., 2023). As a result of this manual extension, overlapping transcript types were resolved based on a hierarchical rule prioritizing genes in the following order: tRNA > mRNA > rRNA > snRNA > snoRNA. The transcript regions (5’ UTR, CDS, intron, 3’ UTR) most frequently observed were assigned to the binding sites to resolve overlapping transcript regions.

To visualize the metaprofile of crosslink events around the stop codon, crosslink events within a 61-nt window centered around the stop codon were extracted from the crosslink coverage for both iCLIP datasets. These crosslink events were normalized using the min-max normalization approach, scaling the minimum value to 0 and the maximum value to 1. This normalization accounted for differences in expression level and other variations. A heatmap of crosslink patterns around the stop codon of specific mitochondrial transcripts was generated using the R Bioconductor package ComplexHeatmap ((Gu et al., 2006), version 2.18.0).

iCLIP2 data are available in the NCBI Gene Expression Omnibus (GEO) under the SuperSeries accession number GSE273496. The security token for anonymous Reviewer access to SuperSeries GSE273496 is xxxxx.

## Acknowledgements

We thank laboratory members and Drs. Heiner Schaal, Alexander Steffen, and Lena Kröninger for critically reading the manuscript. We are grateful to Maria Méndez-Lago and the IMB Genomics Core Facility for their excellent support on iCLIP2 library sequencing and the use of their NextSeq 500 (funded by the Deutsche Forschungsgemeinschaft, DFG, German Research Foundation – P#329045328). In addition, research was funded by grants from the DFG under Germany’s Excellence Strategy EXC-2048/1 – Project ID 39068111 to MF as well as DFG-FOR2333-TP03 to MF, DFG-FOR2333-TP02 to KZ and to JK, Project-ID 267205415 – DFG-SFB 1208 to MF (project A09) as well as Project ID 458090666 – DFG-SFB1535 to MF (project A03).

## Author contributions

NKS, SS, KZ, JK, and MF designed this study and analyzed the data. SS, AB, NKS, and KZ performed the bioinformatic analysis. NKS performed the iCLIP2 experiments supported by NK and JK. NKS, KM, and MF drafted and revised the manuscript with input from all co-authors. MF, ZK, and JK contributed funding and resources.

