## Supplemental Figures 1-5 & Table 1 for "Microbial iCLIP2: Enhanced mapping of RNA-Protein interaction by promoting protein and RNA stability"

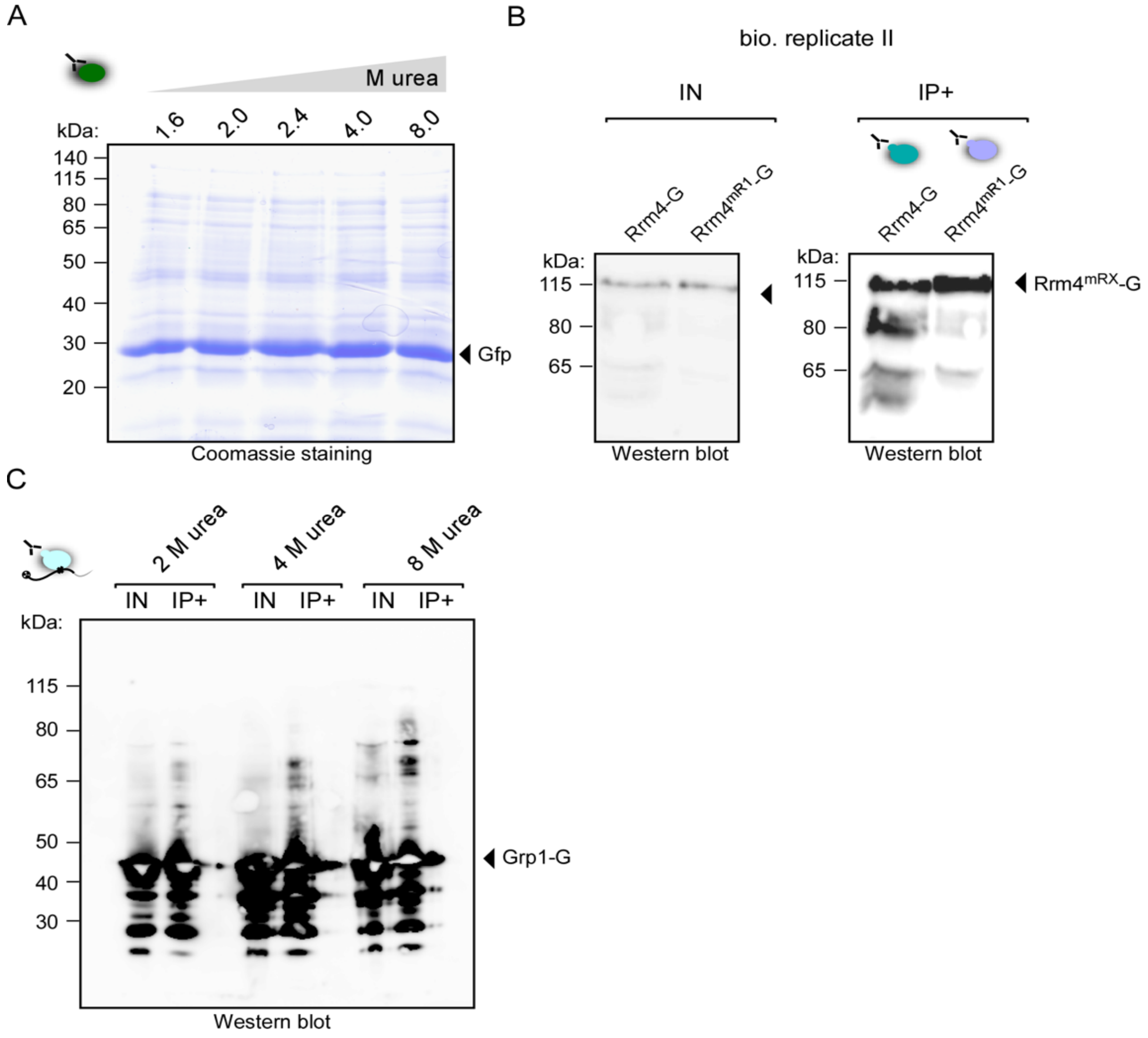


**Supplemental Fig S1. Increased protein stability by improved iCLIP2 protocol. (A)** Coomassie-stained SDS-gel of Gfp IP, purified under different urea molarities conditions. Gfp was expressed in *E.coli* and the respective cell lysate was used as starting material for IP. **(B)** Western Blot analysis Rrm4-Gfp IP (IP+) by the usage of lysing buffer containing 8 M urea. Cell lysate (input = IN) **(C)** Western blot of Grp1-G iCLIP2 (IP+) experiments and respective cell lysate (IN) were produced under different urea conditions (2-8 M urea) - the respective autoradiograph is depicted in Fig. 3A

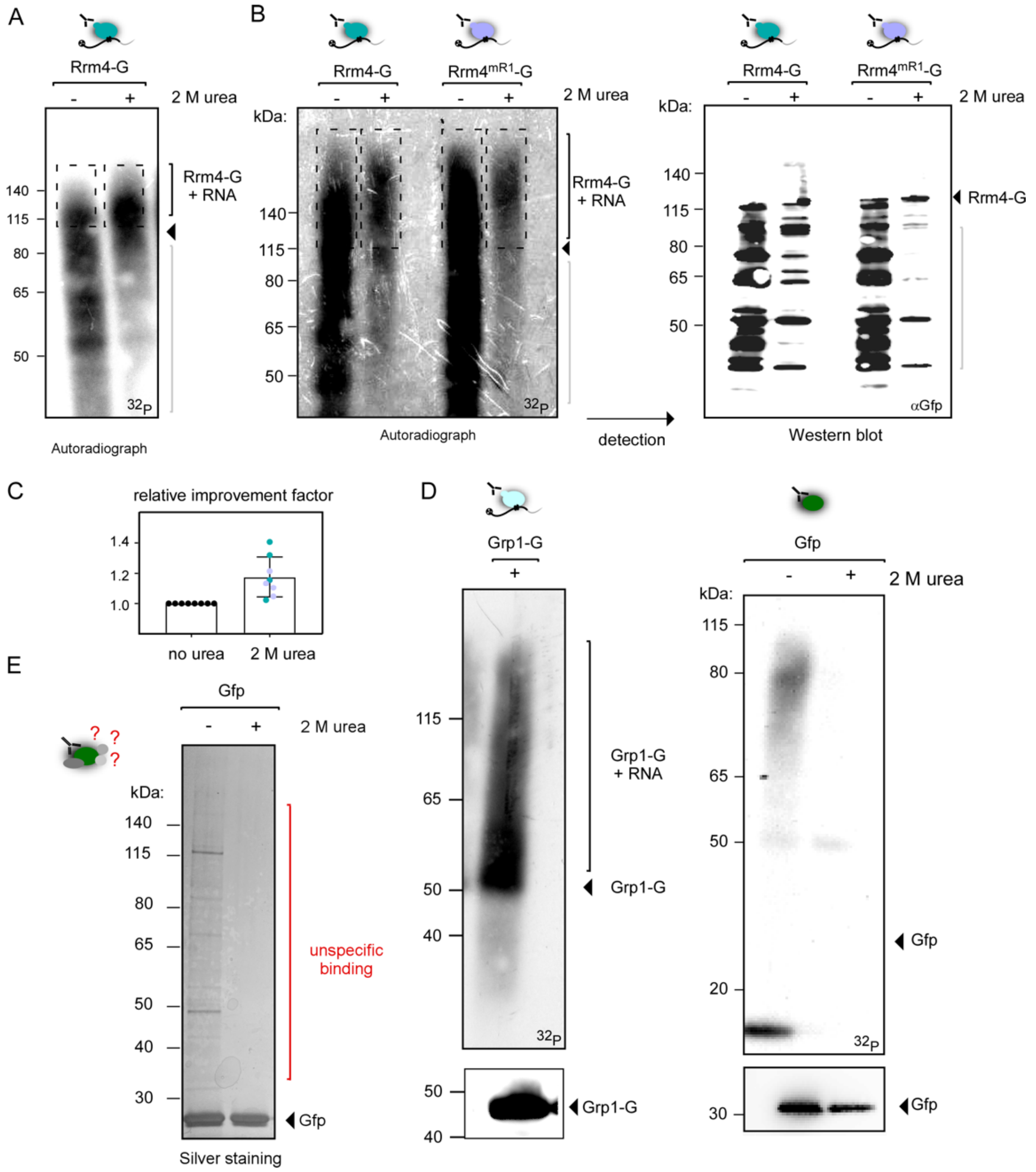


**Supplemental Fig S2. 2 M urea increases protein stability. (A-B)** Autoradiograph of Rrm4-G-RNA complexes by the usage of IPs +/-2 M urea containing lysis buffer (biological replicate). In B the respective Western blot is depicted. **(C)** The relative improvement factor was calculated by dividing the results of relative area quantification (refer to Figure 3C) with the untreated buffer (+ 2M urea/ - urea). **(D)** Autoradiograph of Grp1-G-RNA complexes and free cytoplasmatic Gfp-RNA complexes, purified +/- 2 M urea conditions. Respective Western blots are below. **(E)** Silver-stained SDS-gel from IP Gfp samples prepared -/+ 2 M urea containing lysis buffer. Unspecific co-purified proteins are indicated in red.

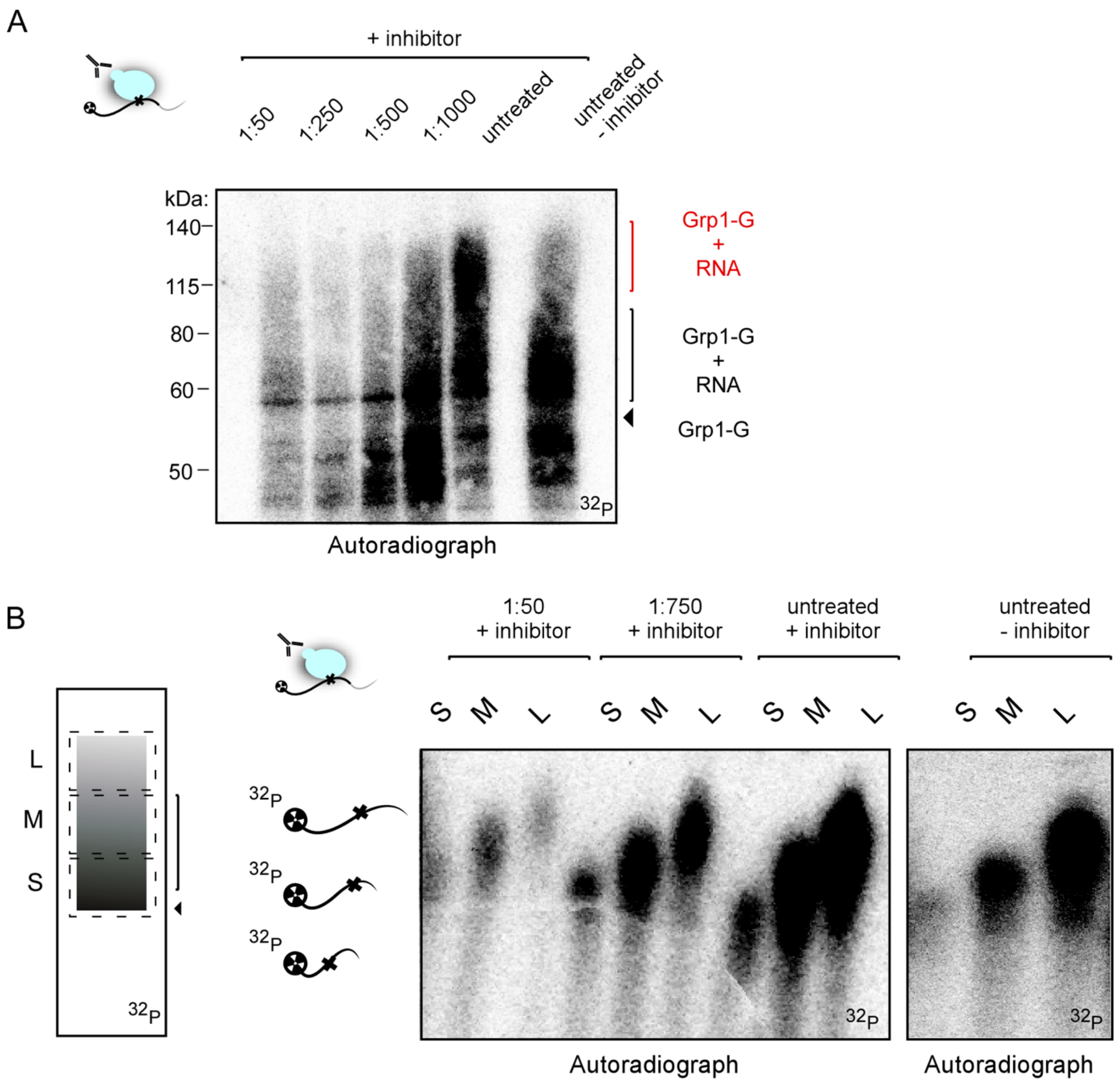


**Supplemental Fig S3. Optimal RNase I condition for microbial iCLIP2 protocol. (A)** Autoradiography of RNase I titration series for Grp1-Gfp. In addition, intrinsic RNase digestion was tested (untreated - inhibitor) The non-adequate-separated Grp1-G-RNA complexes are marked in red. **(B)** Autoradiograph of RNase I titration for Grp1-G-RNA complexes (1:50, 1:750), untreated +/- inhibitor. The respective autoradiograph was separated into short (S), medium (M) and long (L) RNAs. The respective size-selected Grp1-RNA complexes were isolated and separated by a 6 % TBE-Urea-gel as described by Huppertz et al., 2014.

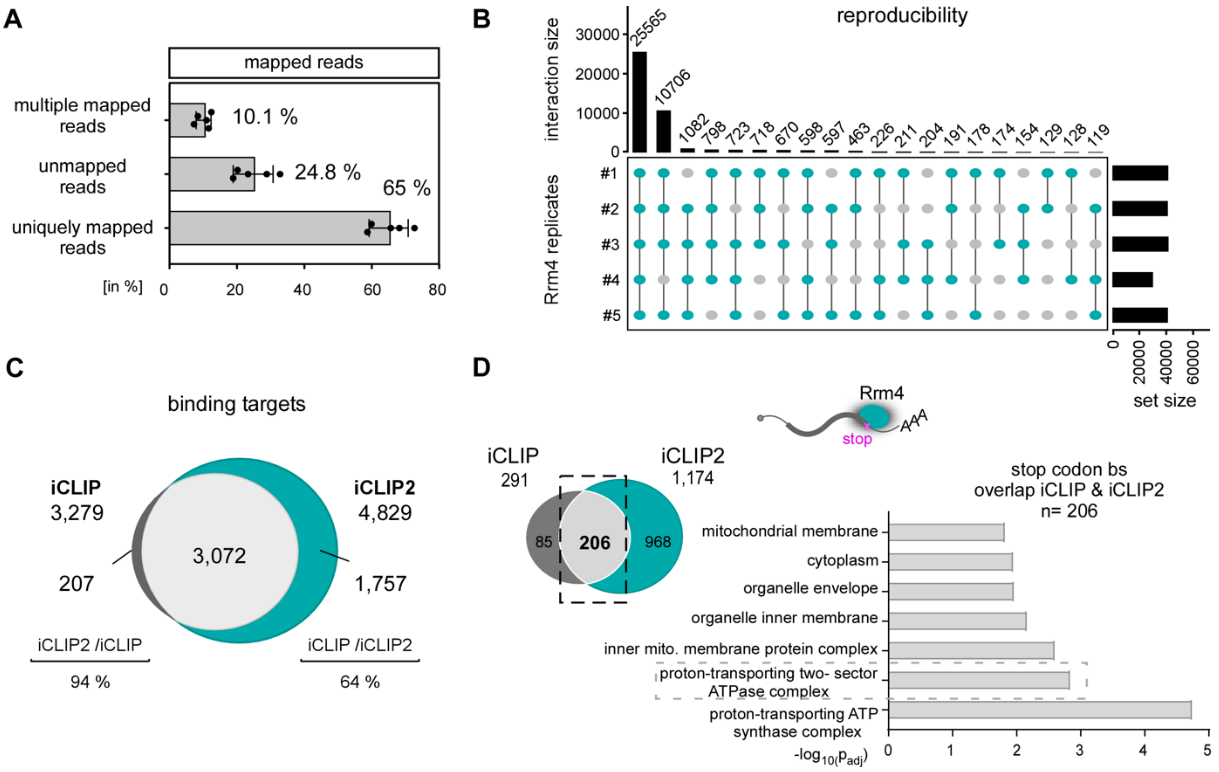


**Supplemental Fig. S4. Improvements achieved in iCLIP2 data set by optimizing the protocol. (A)** Bar chart showing the means of mapped reads from the five biological Rrm4 replicates of iCLIP2. The individual replicates are indicated. The majority of reads are uniquely mapped, as expected. The high number of unmapped reads is due to the low-quality genome of *U. maydis*. Reads within the „unmapped" group for example could be manually determined as rRNAs of *U. maydis*. **(B)** UpSet plot of five biological replicates of iCLIP2 Rrm4. Indicating high reproducibility of the biological replicates **(C)** Venn diagram of iCLIP vs. iCLIP2 Rrm4 overall binding targets **(D)** Bar chart of gProfile analysis of stop codon binding targets determined within both iCLIP and iCLIP2 datasets (overlap n = 206).

**Supplemental Table S1. Summary of different cell lysis conditions**

| Technique | 1<br>Bead mill jar | 2<br>Bead mill tubes | 3<br>Homogenizer | 4<br>Vibrax |
|---|---|---|---|---|
| Cryogenic condition | + | ++ | ++ | |
| Time efficiency | | ++ | ++ | + |
| High throuput/ Volumina | | ++ | ++ | ++ |
| Mechanical separation/ chemical lysis | | ++ | | |

[a] For details of the four different techniques see Materials and methods.
